# PTM-Mamba: A PTM-Aware Protein Language Model with Bidirectional Gated Mamba Blocks

**DOI:** 10.1101/2024.02.28.581983

**Authors:** Zhangzhi Peng, Benjamin Schussheim, Pranam Chatterjee

## Abstract

Proteins serve as the workhorses of living organisms, orchestrating a wide array of vital functions. Post-translational modifications (PTMs) of their amino acids greatly influence the structural and functional diversity of different protein types and uphold proteostasis, allowing cells to swiftly respond to environmental changes and intricately regulate complex biological processes. To this point, efforts to model the complex features of proteins have involved the training of large and expressive protein language models (pLMs) such as ESM-2 and ProtT5, which accurately encode structural, functional, and physicochemical properties of input protein sequences. However, the over 200 million sequences that these pLMs were trained on merely scratch the surface of proteomic diversity, as they neither input nor account for the effects of PTMs. In this work, we fill this major gap in protein sequence modeling by introducing PTM tokens into the pLM training regime. We then leverage recent advancements in structured state space models (SSMs), specifically Mamba, which utilizes efficient hardware-aware primitives to overcome the quadratic time complexities of Transformers. After adding a comprehensive set of PTM tokens to the model vocabulary, we train bidirectional Mamba blocks whose outputs are fused with state-of-the-art ESM-2 embeddings via a novel gating mechanism. We demonstrate that our resultant PTM-aware pLM, **PTM-Mamba**, improves upon ESM-2’s performance on various PTM-specific tasks. PTM-Mamba is the first and only pLM that can uniquely input and represent both wild-type and PTM sequences, motivating downstream modeling and design applications specific to post-translationally modified proteins. To facilitate PTM-aware protein language modeling applications, we have made our model available at: https://huggingface.co/ChatterjeeLab/PTM-Mamba.

## Introduction

Proteins are the key molecular machines of nature. The over 20,000 distinct proteins in the human genome, derived from 20 naturally-occurring amino acids, catalyze diverse biochemical reactions, providing critical scaffolding support to cells, and transporting key molecules throughout organs and tissues [Piovesan et al., 2019]. While the folding of wild-type sequences plays a central role in the function of all proteins, amino acid modifications post-synthesis contribute significantly to the structural and functional diversity of various protein species [Ramazi and Zahiri, 2021, Chen et al., 2023a]. These post-translational modifications (PTMs), including phosphorylation, acetylation, ubiquitination, and glycosylation, together expand the functional capacities of eukaryotic proteomes by various orders of magnitude, as compared to the coding sequence of a genome [Ramazi and Zahiri, 2021, Duan and Walther, 2015]. As such, PTMs influence critical biomolecular processes, including central enzyme activities, protein turnover, and localization dynamics, protein-protein interactions, modulation of unique signaling cascades, cell division, and DNA repair [Ramazi and Zahiri, 2021, Duan and Walther, 2015, Pejaver et al., 2014]. Dysregulation of PTM installation unsurprisingly leads to various disease phenotypes, most notably cancer, neurodegeneration, and aging [Xu et al., 2018, Zhong et al., 2023]. For example, while wild-type STAT3 is a standard transcription factor for eukaryotic cellular function, the phosphorylation of STAT3 at key residues drives the tumorigenesis and metastasis of numerous cancers [Rébé et al., 2013, Lin et al., 2020]. As a result, it is critical to capture the unique features of protein sequences that are post-translationally modified.

The sequential nature of protein sequences, alongside their hierarchical semantics, makes them a natural target for language modeling. Recently, protein language models (pLMs), such as the state-of-the-art ESM-2 and ProtT5 models, have been pre-trained on over 200 million natural protein sequences to generate latent embeddings that accurately encode relevant physicochemical and functional information [Lin et al., 2023, Elnaggar et al., 2022]. Autoregressive pLMs, such as ProGen and ProtGPT2, have generated diverse protein sequences with validated functional capacities [Madani et al., 2023, Ferruz et al., 2022]. Our group has specifically utilized pLM embeddings to *de novo* derive peptides that can bind and degrade diverse target proteins, whether conformationally ordered or not [Palepu et al., 2022, Brixi et al., 2023, Bhat et al., 2023, Chen et al., 2023b]. Arguably the most notable usage of protein sequence embeddings, however, has been for the accurate downstream prediction of protein tertiary structure, as evidenced by the success of models such as AlphaFold2 and ESMFold [Jumper et al., 2021, Lin et al., 2023]. In spite of these incredible advances, *none* of these attention-based models encompass PTM residues as a part of their training or inference procedures, essentially precluding modeling of PTM effects in both sequence and structural space.

In this work, we fill this major gap in protein sequence modeling, by training the first pLM to explicitly encompass PTM information. Instead of using the standard Transformer architecture, we employ recent innovations in state space modeling (SSMs), specifically the Mamba model, which utilizes hardware-aware primitives to overcome the quadratic time complexity of the standard attention mechanism [Gu and Dao, 2023]. For training, rather than relearning the semantics and evolutionary statistics of protein sequences, we curate a comprehensive, experimentally-validated dataset of post-translationally modified protein sequences [UniProt-Consortium, 2020], and leverage a fusion module that dynamically combines the embeddings of PTM-Mamba and ESM-2 in a gated mechanism. Our results demonstrate that by imbuing PTM information into expressive, pre-trained pLM latent spaces, we distinguish the latent representations of PTM sequences from their wild-type counterparts and effectively improve performance on downstream PTM-specific supervised training tasks, including phosphorylation site prediction, non-histone acetylation site prediction, disease association, and druggability. To the best of our knowledge, our resultant **PTM-Mamba** model is the first and only pLM that can input both wild-type and modified protein sequences to derive unique and accurate latent embeddings. As a result, this work motivates the usage of PTM-Mamba to both model and design the complete set of proteoforms found in nature.

## Methods

### Data Curation

Model training data was curated from UniProt [UniProt-Consortium, 2020]. Specifically, 311,350 experimentally-validated PTM records were collected from Swiss-Prot and mapped the PTM annotations of their protein to their respective sequences to construct the new PTM sequences. The final dataset includes a total of 79,707 PTM sequences.

Datasets for the four benchmarking tasks were collected from the following sources. Phosphorylation site data was obtained from the corresponding ProteinBERT benchmark [Brandes et al., 2022], originally derived from PhospoSitePlus [Hornbeck et al., 2011], and filtered for sequences between 256 and 512 amino acids in length, yielding a training set of 15,588 sequences, a validation set of 1707 sequences, and a testing set of 3106 sequences. Non-histone acetylation site prediction was performed equivalently as described in prior literature, using the NHAC (Non-Histone Acetylation Collection) dataset [Meng et al., 2023]. The druggability and disease association datasets were curated from the dbPTM database [Li et al., 2021]. Briefly, wild-type sequences were mapped to corresponding entries in the PTM-Mamba dataset, and wild-type residues were replaced by the corresponding, position-specific PTMs for either PTM-Mamba tokenization or one-hot embeddings. For all other baseline models trained with standard one-hot embeddings or ESM-2 embeddings, the corresponding wild-type sequence was used as input.

### Tokenization

In our tokenization scheme, we use the standard set of amino acids tokens as described in ESM-2 [Lin et al., 2023]. In addition to special tokens, the 20 wild-type amino acids tokens are as follows: D, N, E, K, V, Y, A, Q, M, I, T, L, R, F, G, C, S, P, H, W. We introduce new PTM tokens, corresponding to their unique specific UniProt annotations: <N-linked (GlcNAc…) asparagine>, <Pyrrolidone carboxylic acid>, <Phosphoserine>, <Phosphothreonine>, <N-acetylalanine>, <N-acetylmethionine>, <N6-acetyllysine>, <Phosphotyrosine>, <S-diacylglycerol cysteine>, <N6-(pyridoxal phosphate)lysine>, <N-acetylserine>, <N6-carboxylysine>, <N6-succinyllysine>, <S-palmitoyl cysteine>, <O-(pantetheine 4-phosphoryl)serine>, <Sulfotyrosine>, <O-linked (GalNAc…) threonine>, <Omega-N-methylarginine>, <N-myristoyl glycine>, <4-hydroxyproline>, <Asymmetric dimethylarginine>, <N5-methylglutamine>, <4-aspartylphosphate>, <S-geranylgeranyl cysteine>, <4-carboxyglutamate>. The top two most abundant PTM tokens are: `<N-linked (GlcNAc…) asparagine>` and `<Phosphoserine>`. The full distribution of the PTM tokens is shown in Supplementary Figure 1. The wild-type amino acid tokens are then converted into embeddings by both ESM-2-650M and PTM-Mamba, while the PTM tokens are only processed by PTM-Mamba.

### PTM-Mamba Architecture

PTM-Mamba is a bidirectional pLM that represents both amino acids and PTM tokens. The model builds on top of the state-of-the-art, structured SSM, Mamba [Gu and Dao, 2023], with an added bidirectional module and gated embedding fusion module. Briefly, Mamba is a novel class of structured SSM, an alternative to convolutional neural networks (CNNs), recurrent neural networks (RNNs), or Transformers, that model the long-range sequential dependency with sub-quadratic complexity. Despite its effectiveness in sequential modeling, Mamba was designed for autoregressive text generation, thus not capturing full sequence semantics. Here, we build upon the standard Mamba block (Figure 1A) introduced previously [Gu and Dao, 2023] to propose a unique bidirectional Mamba block (Figure 1B and Listing 1), consisting of forward and backward Mamba blocks which individually process the sequences from left to right and right to left, respectively.

**Figure 1:**
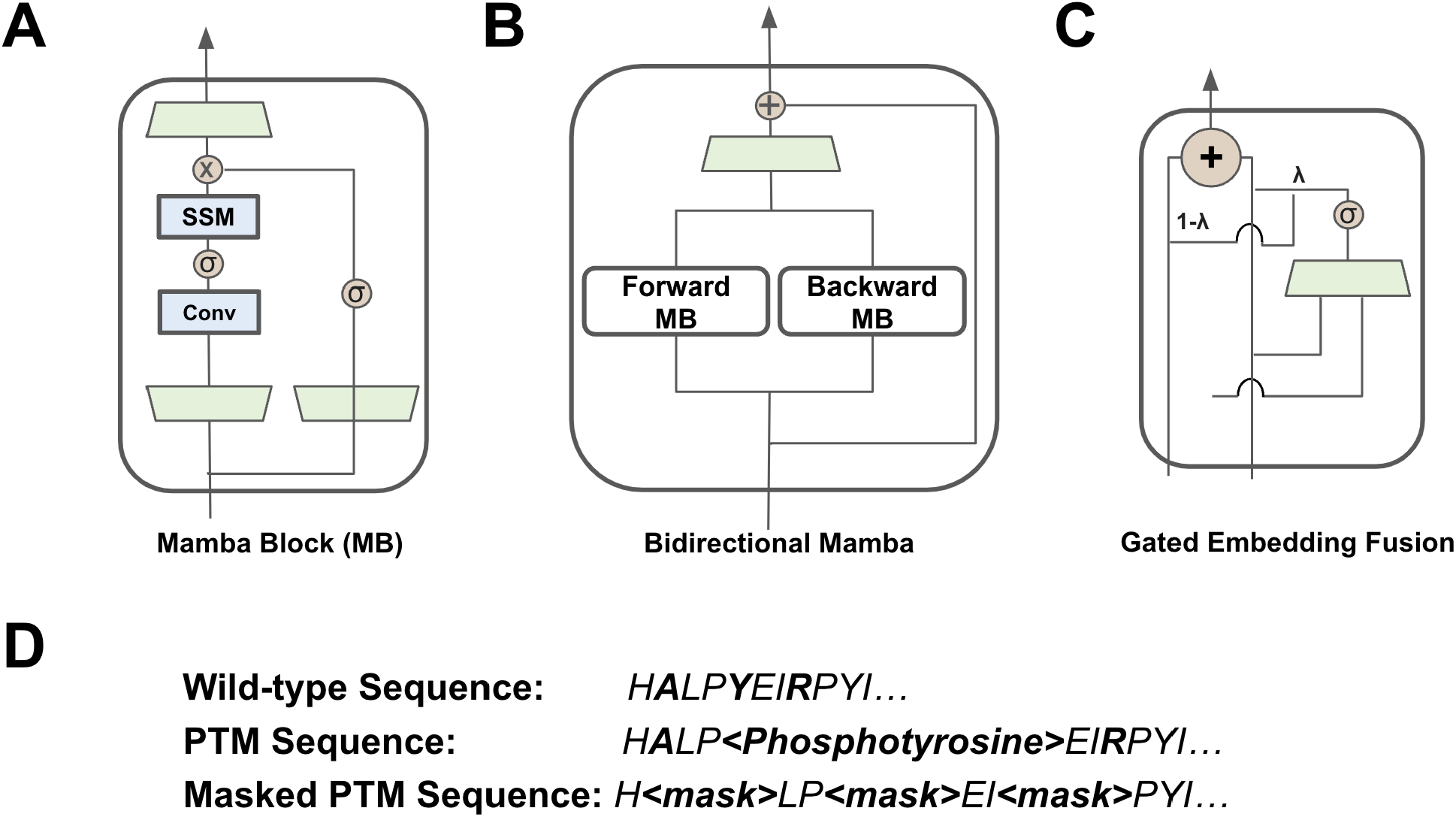
Primitives of PTM-Mamba. A) The standard building block of Mamba [Gu and Dao, 2023]. B) The bidirectional Mamba block in PTM-Mamba is built on top of the Mamba block in (A), which processes the sequences forward (Forward MB) and backward (Backward MB). C) The gated embedding fusion module inputs the ESM-2 and PTM embeddings and fuses them in a gated manner via a sigmoid-activated linear layer. D) Wild-type and PTM sequences, where some residues are replaced with their PTM annotations. Masked sequences are obtained by randomly masking the PTM sequence for training.

**Listing 1:**
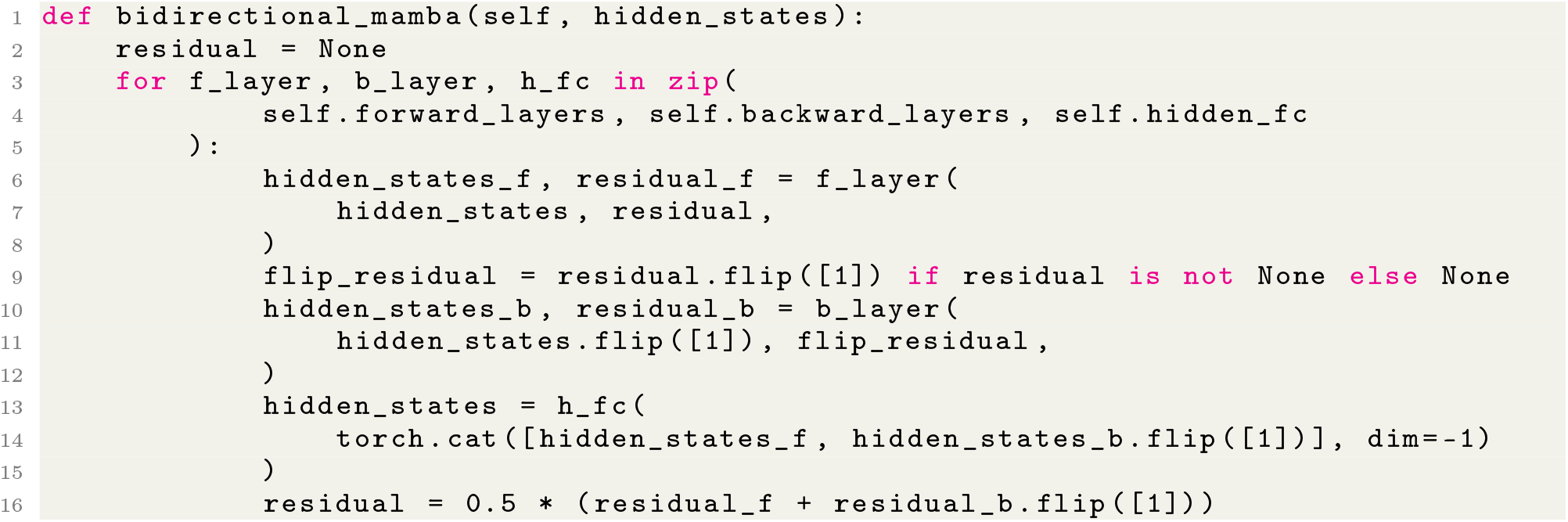
Bidirectional Mamba block implementation.

To preserve comprehension of regular amino acids, we train PTM-Mamba as a head to the state-of-the-art ESM-2-650M model, where wild-type amino acid tokens are passed into ESM-2-650M to retrieve its output embeddings and PTM tokens are converted into `<mask>` tokens for ESM-2-650M input. Sequences are finally fed into the embedding layer of the PTM-Mamba, which naturally processes both wild-type and PTM tokens. To join the ESM-2-650M and PTM-Mamba embeddings, we propose a novel gating mechanism, where the two embeddings are concatenated and filtered via a sigmoid-activated linear gate to produce a final output representation (Figure 1C and Listing 2).

**Listing 2:**
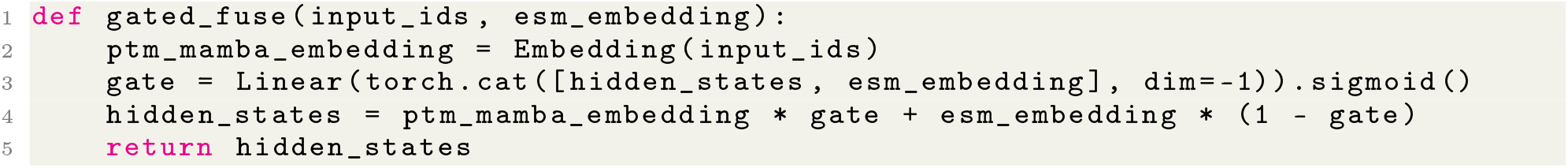
Gated Embedding Fusion Module.

### PTM-Mamba Training Procedure

PTM-Mamba was trained on a Nvidia 8xA100 DGX system with 640 GB of shared VRAM on an adjusted masked language modeling (MLM) task, where rather than random 15% token masking, we bias masking to PTM residue tokens (Figure 1D). Briefly, given a sequence, with 80% probability, we perform standard 15% token masking, and with 20% probability, we mask all the PTM tokens and randomly mask 15% of wild-type tokens. For training, we then consider a protein sequence with masked residues, where the model aims to predict the original tokens at these residue positions. Let *x*_*i*_ denote the original residue token at position *i* that has been masked, and *y*_*i*_ denote the residue token predicted by the model for this position. The loss function *L* for MLM can be defined as the negative log-likelihood of the correct tokens given their masked inputs, summed over all masked positions *N* :

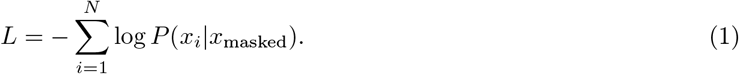

*P* (*x*_*i*_|*x*_masked_) represents the probability of predicting the correct original token *x*_*i*_ at the masked position, given the masked input sequence *x*_masked_. PTM-Mamba was trained via the `Adam` optimizer with no weight decay. The final PTM-Mamba model has `24` layers with hidden dimensions of `768`. It was trained for 400K steps at a constant learning rate of `0.0002` with batch size `256` and dynamic batching. Training sequences were randomly cropped to a maximal length of 1024 or padded at the end to reach a length of 1024. During training, we clustered the sequences by length and constructed the batches. The training batches were fed into the model, going from the shortest to the longest sequences.

### Benchmark Model Training

For phosphorylation site prediction, models employing baseline embeddings and PTM-Mamba embeddings were trained via multilayer perceptron (MLP) with a tanh-activated `Linear` hidden layer and two `Dropout` layers. These models were trained in PyTorch using an `Adam` optimizer with a learning rate of `lr = 0.001` using a weight-balance of `[0.01, 0.99]`. Employing the ESM-2-650M and PTM-Mamba embeddings, non-histone acetylation site prediction was performed as described in prior literature, using the equivalent model architecture and train/test dataset [Meng et al., 2023]. The disease association and druggability models were trained via a one-layer MLP model in PyTorch, using an `Adam` optimizer with a learning rate of `lr = 0.001`. For models trained on one-hot embeddings of wild-type input sequences, an `nn.Embedding` layer with a hidden dimension of 1280 was utilized. Models were all trained for 40 epochs using one Nvidia A100 GPU with 80 GB of VRAM. Models were evaluated using the Accuracy, Precision, Recall, F1 Score, MCC, AUROC, and AUPRC metrics via `scikit-learn`.

## Results

### PTM-Aware Embedding Exploration

The primary objective of PTM-Mamba is to distinctly, yet relevantly, represent both unmodified and post-translationally modified protein sequences, acknowledging the critical biological functions and structural changes induced by PTMs [Ramazi and Zahiri, 2021]. Given this aim, we visualize PTM-Mamba embeddings in a two-dimensional context to concretely assess the model’s capability in achieving embedding differentiation. Via t-SNE visualization of the generated embeddings, we indeed observe a nuanced distinction between embeddings of wild-type protein sequences and that of their PTM counterparts (Figure 2A). The proximity of the embeddings for each wild-type: PTM-modified pair indicates that PTM-Mamba successfully captures the subtle, yet important, differences imparted by PTMs. This proximity reflects the model’s ability to recognize the semantic relevance of PTMs while maintaining the contextual integrity of the protein sequence. Our observations are further supported by the unique token embeddings generated by PTM-Mamba for PTM tokens vs. wild-type amino acid tokens (Figure 2B). As validation of successful token embedding generation, we note that modified residue tokens belonging to specific PTM classes, such as phosphorylation and acetylation, are spatially proximal (Figure 2B). Furthermore, based on token embedding distribution, it is clear that PTM-Mamba strongly attends to PTM residue tokens, compared to less spatially diverse wild-type tokens, indicating that the model focuses effort on augmenting ESM-2 embeddings with PTM residue information (Figure 2B).

**Figure 2:**
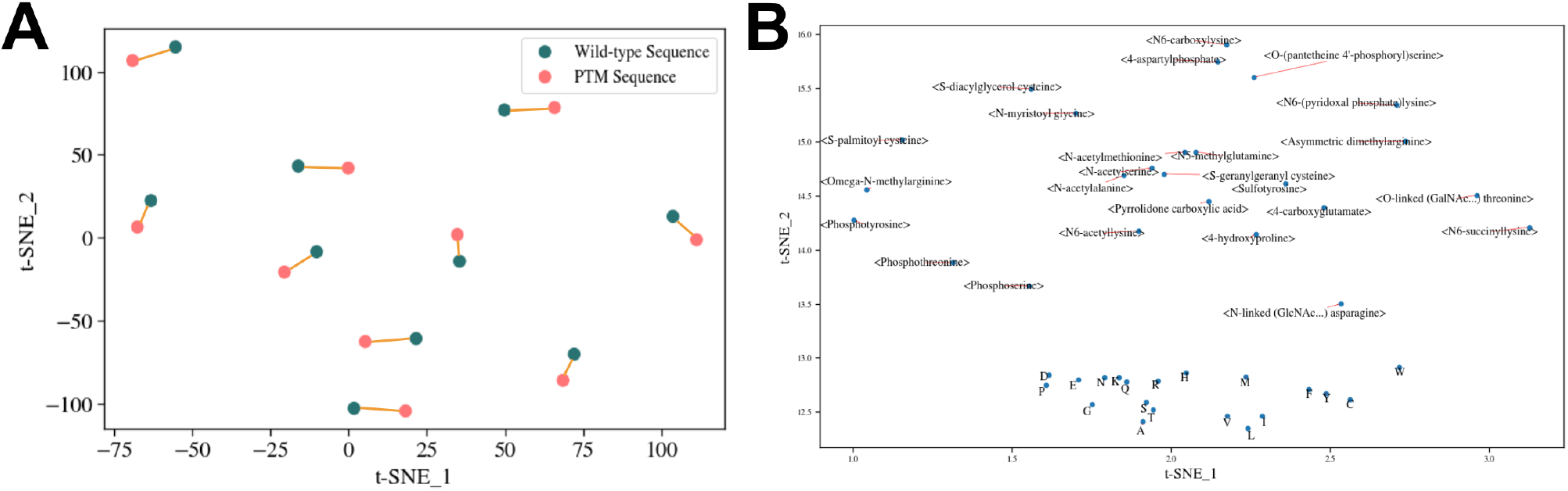
Visualization of PTM-Mamba sequence and token embeddings. A) t-SNE visualization of embeddings of example wild-type and corresponding PTM sequences. Orange lines connect the corresponding embeddings. B) t-SNE visualization of labeled token embeddings.

### Phosphorylation Site Prediction

One of the most well-characterized and consequential PTMs is phosphorylation, which can mediate cell growth, differentiation, and apoptotic pathways [Ramazi and Zahiri, 2021, Ardito et al., 2017]. To evaluate the quality of PTM-Mamba embeddings on downstream prediction tasks, we first focused on predicting phosphorylation sites within input protein sequences. From a previous pLM study [Brandes et al., 2022, Hornbeck et al., 2011], we curated a dataset of phosphorylation site annotations, and conducted per-residue binary classifications for input sequences. As a clear demonstration of its expressive PTM-aware latent representations, over all classification metrics, PTM-Mamba strongly outperforms all other models, including ESM-2-650M (Figure 3A). The superior performance of PTM-Mamba embeddings is maintained regardless of sequence length, underscoring its generalizability to various protein species (Figure 3B-D and Supplementary Figure 2). Moreover, we note that while PTM-Mamba preferably attends to PTM residue tokens (Figure 2B), this benchmark dataset only contains wild-type amino acid sequences, suggesting that PTM-Mamba can be used in lieu of ESM-2 for PTM-specific property prediction of wild-type sequences as well.

**Figure 3:**
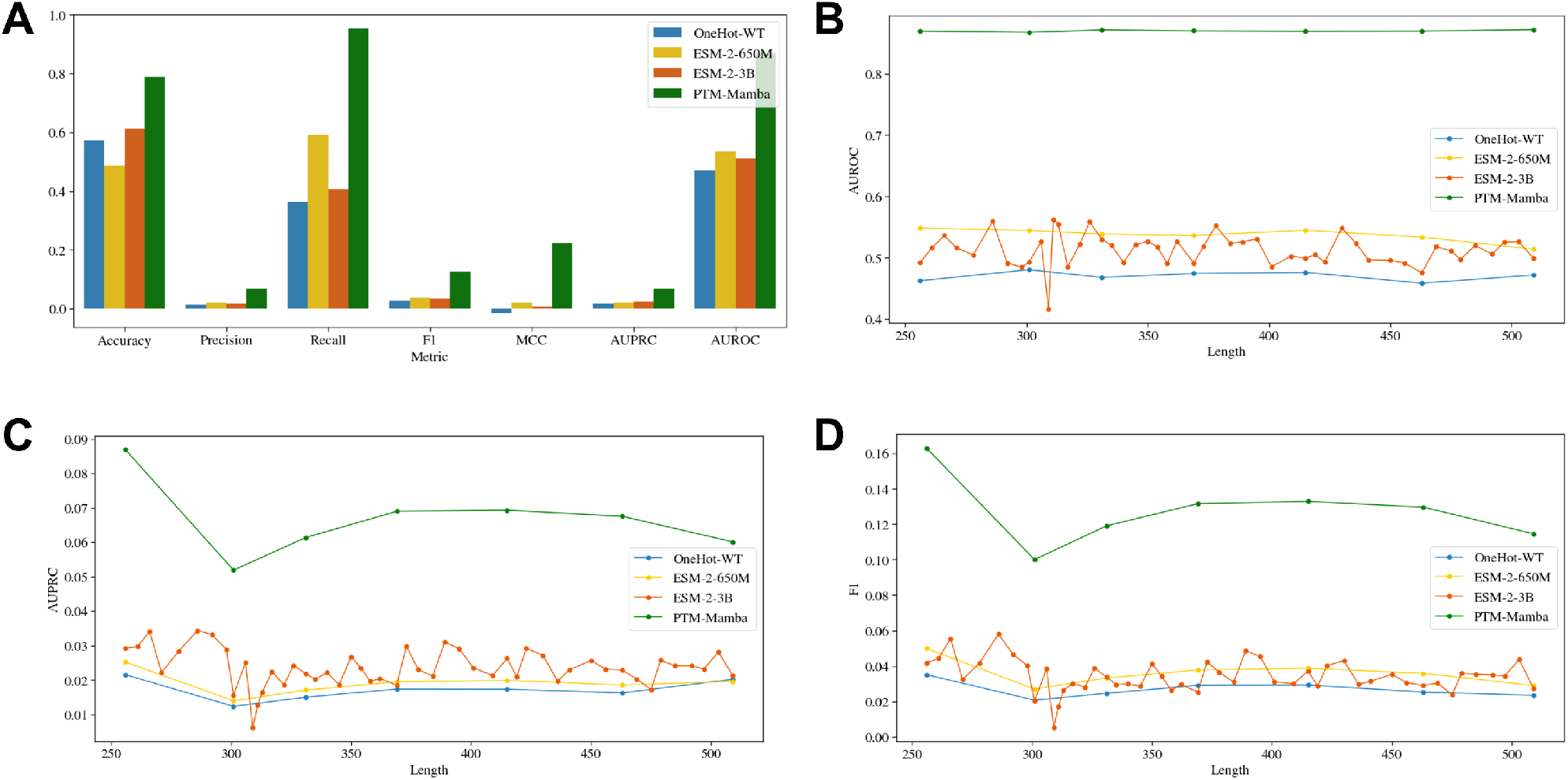
Evaluation of phosphorylation site prediction. A) Classification metrics of baseline models vs. PTM-Mamba. B) AUROC of site prediction for different length input protein sequences. C) AUPRC of site prediction for different length input protein sequences. D) F1 score of site prediction for different length input protein sequences.

### Non-Histone Acetylation Site Prediction

We next turned our attention to protein acetylation, a PTM extensively explored for its pivotal roles in gene regulation, protein stability, signal transduction, cellular metabolism, apoptosis, and DNA repair [Narita et al., 2018]. Specifically, we sought to evaluate PTM-Mamba’s ability to predict non-histone acetylation sites as compared to ESM-2-650M, by training a per-residue binary classifier on the recently-curated Non-Histone Acetylation Collection (NHAC) dataset [Meng et al., 2023]. Our results demonstrate that PTM-Mamba outperforms ESM-2-650M on all classification metrics over the entire test dataset (Figure 4A), including at distinct input sequence lengths (Figure 4B-D and Supplementary Figure 3), highlighting the added benefit of PTM representations to ESM-2-650M embeddings and motivating usage of PTM-Mamba for additional PTM site prediction tasks.

**Figure 4:**
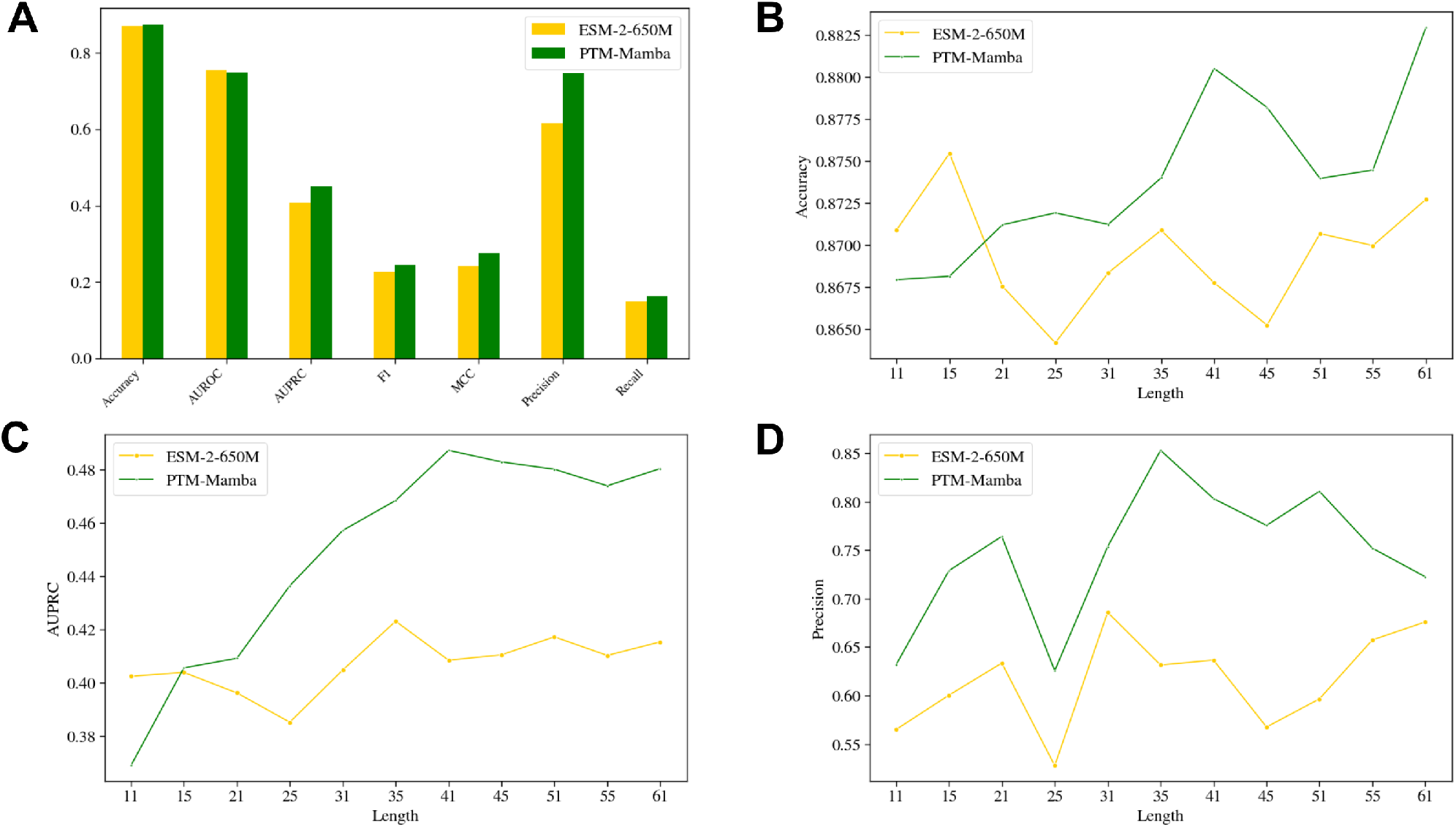
Evaluation of non-histone acetylation site prediction. A) Classification metrics of ESM-2-650M vs. PTM-Mamba. B) Accuracy of site prediction for different length input protein sequences. C) AUPRC of site prediction for different length input protein sequences. D) Precision metric of site prediction for different length input protein sequences.

### Disease Association Prediction of Modified Sequences

While PTM installation on wild-type protein sequences is critical for regulating protein function and cellular signaling pathways, incorrect or aberrant PTMs can lead to diseases by altering these regulatory mechanisms. In cancer, the methylation of arginine 17 on histone H3 (H3R17me), for example, can alter chromatin structure and gene expression, promoting oncogene activation and tumor progression [Poulard et al., 2016]. The abnormal phosphorylation of tau protein at threonine 231 (tau-pT231) is a key event in Alzheimer’s disease, leading to tau dysfunction, neurofibrillary tangle formation, and neuronal death, contributing to the disease’s progression [Ashton et al., 2022]. As such, we sought to evaluate whether the utilization of PTM-Mamba’s unique PTM token embeddings could improve the disease association classification of modified sequences. After curating a comprehensive labeled dataset of PTM sequences for this task from dbPTM [Li et al., 2021], we trained a simple MLP classifier on baseline embeddings, including dummy one-hot embeddings either including or excluding PTMs, as well as the ESM-2-650M and ESM-2-3B models, which do not explicitly tokenize PTM residues. While minimal differences were observed between one-hot embeddings across relevant classification metrics, ESM-2-650M outperformed ESM-2-3B on this task, justifying its usage within PTM-Mamba. Most importantly, PTM-Mamba was the clear top performer on the relevant metrics, including AUROC and AUPRC, demonstrating the utility of its PTM token representations for capturing PTM-specific effects (Table 1 and Supplementary Figure 4A-B).

**Table 1:** Evaluation of PTM-Mamba vs. baseline models on disease association of PTM sequences.

| Model | Accuracy | Precision | Recall | F1 Score | MCC | AUROC | AUPRC |
| --- | --- | --- | --- | --- | --- | --- | --- |
| OneHot-WT | 0.657 | 0.653 | **0.990** | 0.786 | 0.135 | 0.722 | 0.832 |
| OneHot-PTM | 0.656 | 0.652 | **0.990** | 0.786 | 0.13 | 0.722 | 0.832 |
| ESM-2-650M | 0.662 | 0.667 | 0.936 | 0.78 | 0.178 | 0.737 | 0.836 |
| ESM-2-3B | 0.593 | 0.681 | 0.676 | 0.68 | 0.122 | 0.531 | 0.656 |
| PTM-Mamba | **0.692** | **0.696** | 0.914 | **0.792** | **0.276** | **0.753** | **0.847** |

### Druggability Prediction of Modified Sequences

PTMs can significantly influence the druggability of proteins by altering their structure, function, and interaction networks, thereby impacting the efficacy of potential therapeutic agents [Zhong et al., 2023]. For example, PTMs such as phosphorylation or ubiquitination can change the conformation of a protein, potentially revealing or obscuring binding sites for small molecule drugs, thereby affecting a drug’s ability to modulate the protein’s activity in disease contexts [Lee et al., 2023]. By leveraging a pre-processed labeled dataset that labels druggable PTM sequences [Li et al., 2021], we next decided to benchmark PTM-Mamba against the aforementioned baseline embeddings on druggability classification. For negative labels, we isolated protein sequences from a dataset of undruggable cancer targets [Behan et al., 2019], and their associated Swiss-Prot PTM annotations. After training a simple MLP classifier on baseline embeddings and PTM-Mamba embeddings, we demonstrate that PTM-Mamba exhibits improved performance over ESM-2-based and one-hot-based models on the critical Accuracy, Precision, F1 score, and MCC metrics while exhibiting similar performance to ESM-2-650M on AUROC and AUPRC (Table 2 and Supplementary Figure 4C-D). These results both highlight the utility of base ESM-2 wild-type embeddings and demonstrate the added benefit of PTM-aware embeddings for downstream property prediction tasks.

**Table 2:** Evaluation of PTM-Mamba vs. baseline models on the druggability of PTM sequences.

| Model | Accuracy | Precision | Recall | F1 Score | MCC | AUROC | AUPRC |
| --- | --- | --- | --- | --- | --- | --- | --- |
| OneHot-WT | 0.767 | 0.777 | **0.978** | 0.866 | 0.067 | 0.686 | 0.84 |
| OneHot-PTM | 0.767 | 0.777 | **0.978** | 0.866 | 0.067 | 0.687 | 0.84 |
| ESM-2-650M | 0.855 | 0.858 | 0.975 | 0.912 | 0.55 | **0.921** | **0.976** |
| ESM-2-3B | 0.835 | 0.878 | 0.879 | 0.891 | 0.545 | 0.851 | 0.942 |
| PTM-Mamba | **0.861** | **0.886** | 0.947 | **0.913** | **0.581** | 0.905 | 0.967 |

### Zero-Shot PTM Discovery

Finally, to assess the functional plausibility of PTM-Mamba embeddings in a zero-shot fashion, we analyzed the model’s output logits for specified amino acids in randomly downselected wild-type sequences. For these sequences, we mask each position and compute the logits of the masked positions. We show that the largest logits correspond to PTMs that are capable of being installed on the wild-type amino acid (Figure 5). As an example, for the serine residue at position 89 in Q02261, the largest output logit value corresponds to `<Phosphoserine>` and for cysteine 20 in Q4L7X2, the preferred PTM token is `<S-diacylglycerol cysteine>` (Figure 5). These results underscore the effectiveness of PTM-Mamba as a potential PTM discovery tool, showcasing the model’s potential for capturing nuanced biological insights.

**Figure 5:**
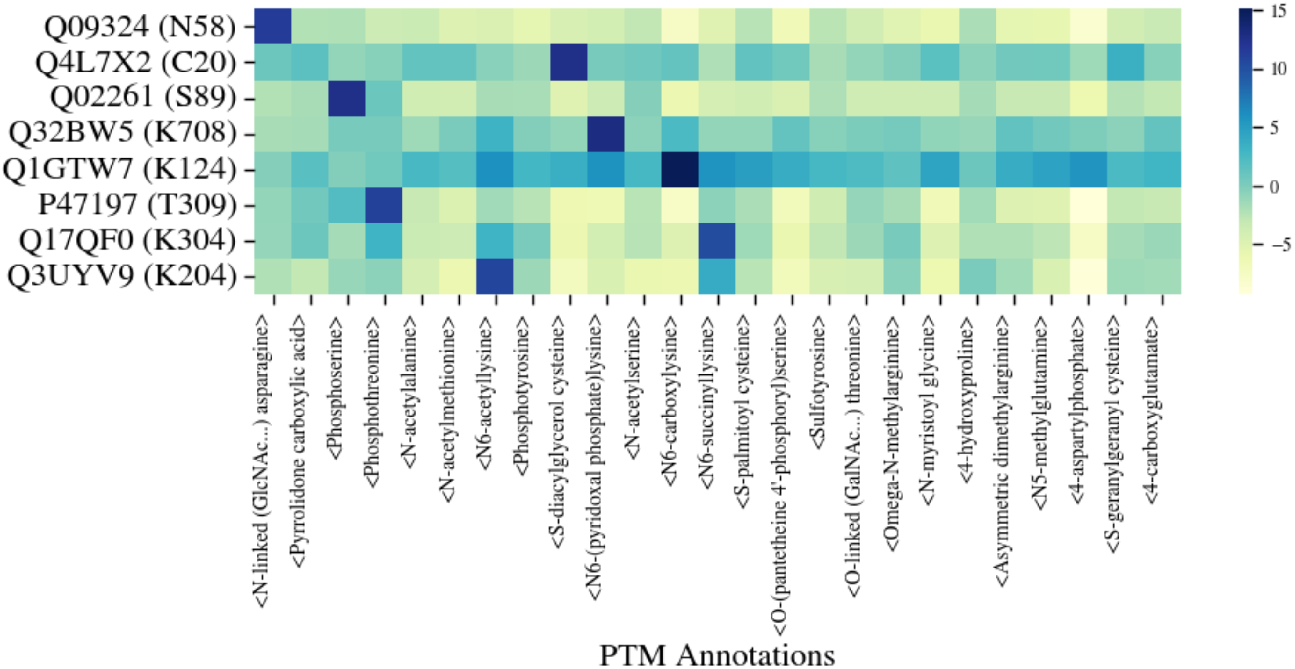
Visualization of the predicted logits for zero-shot PTM discovery. The row denotes different amino acids in the format of `Uniref-accession-id (amino-acid position)` and the columns denote the logit value of the PTMs.

## Discussion

In this work, we introduce the first pLM trained to represent both wild-type and post-translationally modified protein sequences, capturing PTM-specific effects in its latent space. To do this, we trained a bidirectional gated Mamba block to integrate ESM-2 embeddings with new PTM residue tokens, enabling augmentation of ESM-2’s expressive latent space to modified protein sequences. To our knowledge, PTM-Mamba is the only model that enables input of protein sequences with PTM-altered residues specifically encoded at inference time. Architecturally, we further contribute a novel gating strategy to integrate new token embeddings into pre-trained model architectures. This work, overall, highlights the power of SSM primitives for protein language modeling, representing one of the first use cases of these subquadratic architectures on protein sequences [Zhang and Okumura, 2024], and the first use case of the powerful, hardware-efficient Mamba architecture for protein modeling [Gu and Dao, 2023].

We do note, however, that future revisions to both our model and evaluation criteria will improve the utility of PTM-Mamba. First and foremost, PTM-Mamba suffers from both long-tail distribution of PTM labels and data scarcity, as it is trained on around 80,000 PTM sequences from the Swiss-Prot database with known PTM annotations (Supplementary Figure 1). In future work, we can augment training data with unreviewed TrEMBL entries, which may contain additional predicted annotations of lower confidence proteins that can prove useful for training [UniProt-Consortium, 2020]. Furthermore, while we demonstrate PTM-Mamba’s improved relative performance in predicting disease association and druggability of PTM sequences, these labels represent higher-order, more generalized properties of protein sequences [Li et al., 2021]. In addition to validating PTM-Mamba embeddings on standard, wild-type benchmarking tasks [Dallago et al., 2021, Notin et al., 2023], we plan to curate datasets of PTM-specific biophysical labels, including annotated binding sites, cellular localization, and protein stability of PTM sequences. Additional benchmarks leveraging these datasets will provide improved confidence in using PTM-aware embeddings for downstream modeling tasks.

Nonetheless, our current PTM-Mamba pLM presents exciting opportunities for sequence-based protein design. Building off of our previous work in designing protein-binding peptides provided only the target sequence [Bhat et al., 2023, Chen et al., 2023b], specifically enabling binder design to conformationally-disordered target proteins, we envision that a PTM-aware pLM will enable the design of binders that recognize specific post-translational protein states. As such, next steps include augmenting our PepMLM architecture, which leverages a novel peptide masking strategy on ESM-2 embeddings [Chen et al., 2023b], with PTM-Mamba embeddings, enabling binder design to specific modified proteoforms. Our ultimate goal will be to leverage PTM-Mamba to computationally design and experimentally validate clinically-viable biologics that selectively modulate modified disease-causing proteins versus their wild-type healthy counterparts. As such, our platform, now incorporating PTM information, will impart a level of targeting specificity that is currently intractable within standard drug discovery pipelines.

## Declarations

## Acknowledgements

We thank Mark III Systems for computing support. We further thank Sophia Vincoff, Yinuo Zhang, and Tianlai Chen for their insights related to the manuscript. The work was supported by the National Cancer Institute (Award #R21CA278468).

## Author Contributions

Z.P. designed and implemented PTM-Mamba architecture. Z.P. and B.S. performed benchmarking and analysis. Z.P. and P.C. wrote and reviewed the manuscript. P.C. conceived, designed, directed, and supervised the study.

## Data and Materials Availability

All data needed to evaluate the conclusions are presented in the paper and tables. Model weights and training code can be found at: https://huggingface.co/ChatterjeeLab/PTM-Mamba and https://github.com/programmablebio/ptm-mamba.

## Supplementary Material

**Supplementary Figure 1:**
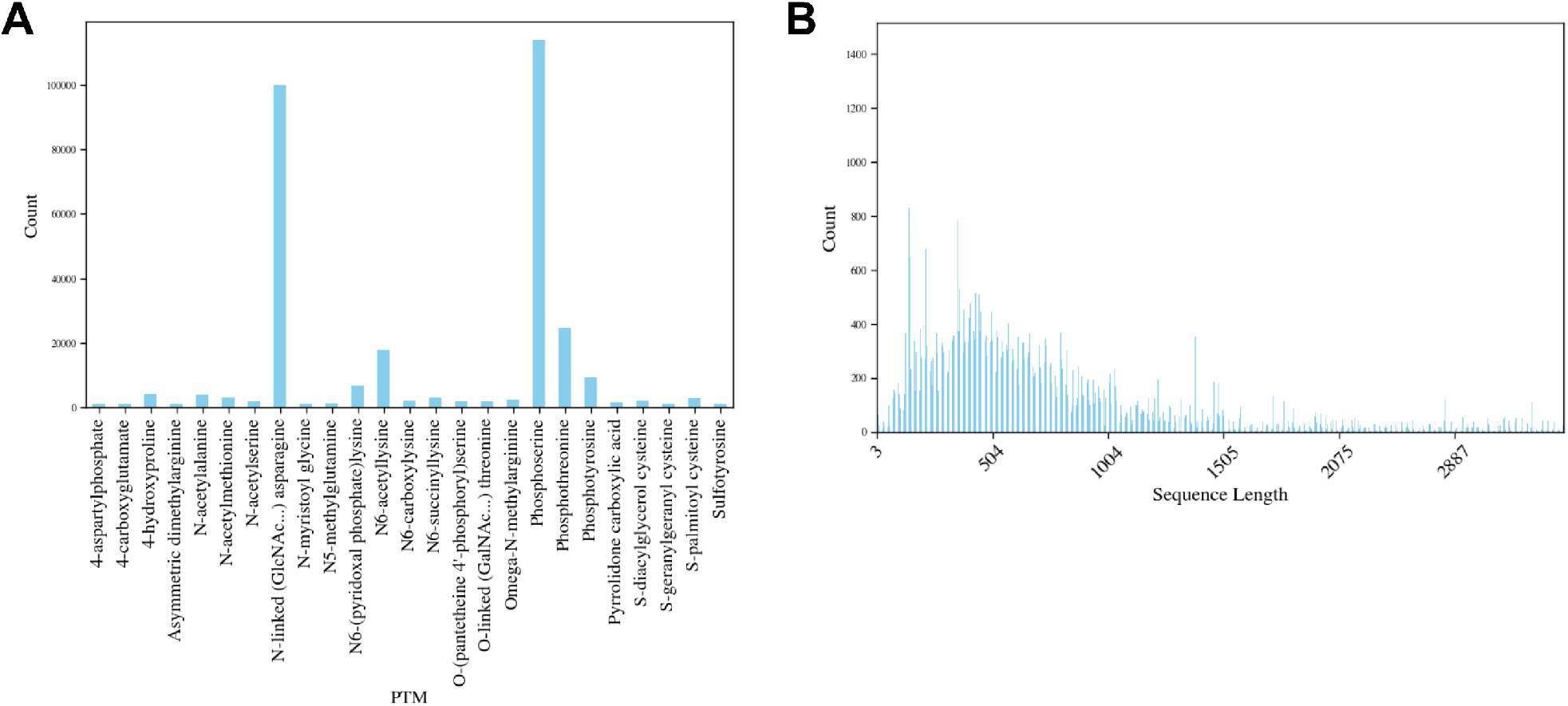
PTM distributions. A) The distribution of PTM annotations, demonstrating long-tail distributions. B) The distribution of the PTM sequence length, which is heavily centered at 400. A majority of PTM sequences are below length 1500.

**Supplementary Figure 2:**
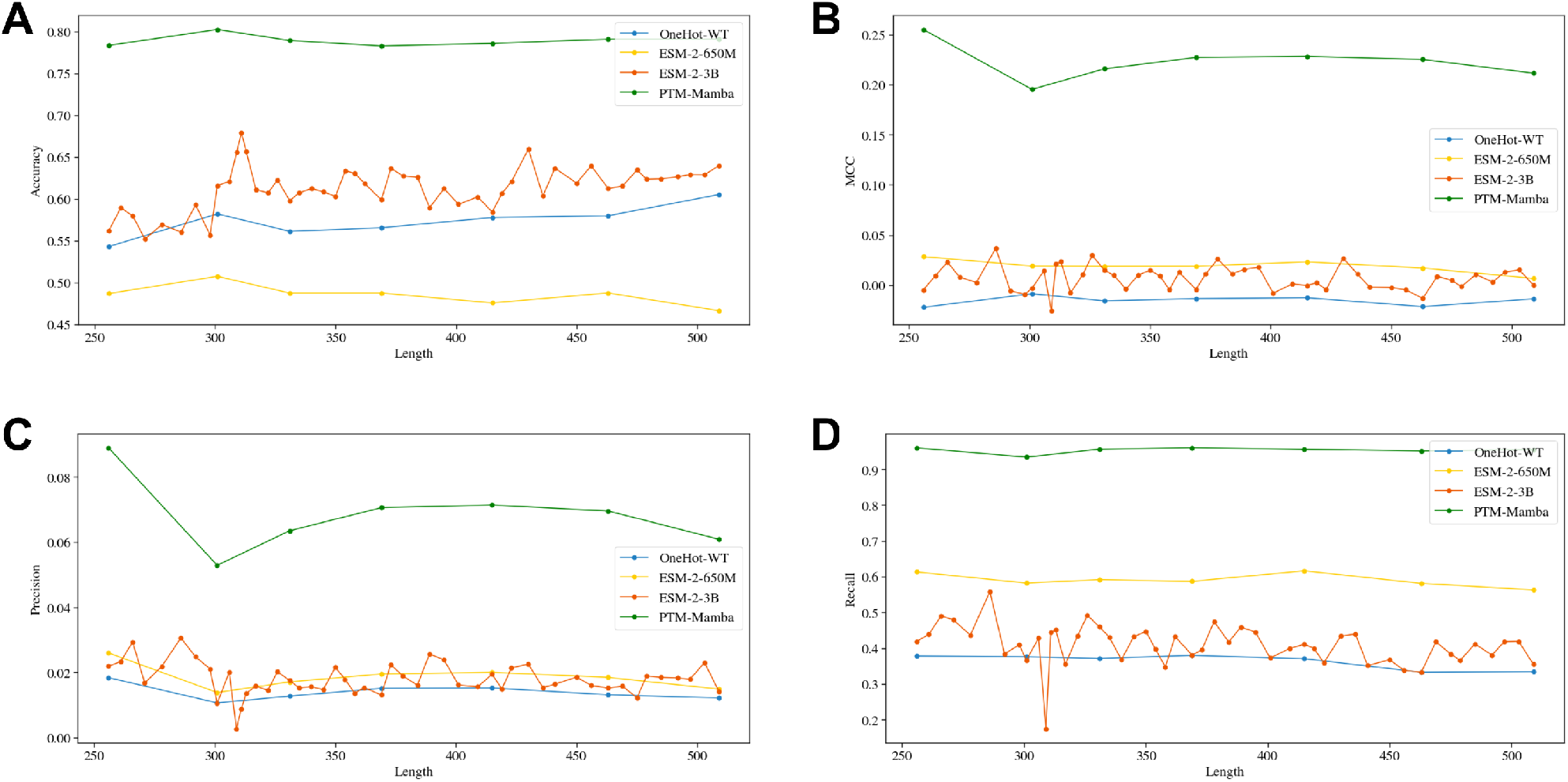
Alternate metrics of phosphorylation site prediction. A) Accuracy of site prediction for different length input protein sequences. B) MCC metric of site prediction for different length input protein sequences. C) Precision of site prediction for different length input protein sequences. D) Recall of site prediction for different length input protein sequences.

**Supplementary Figure 3:**
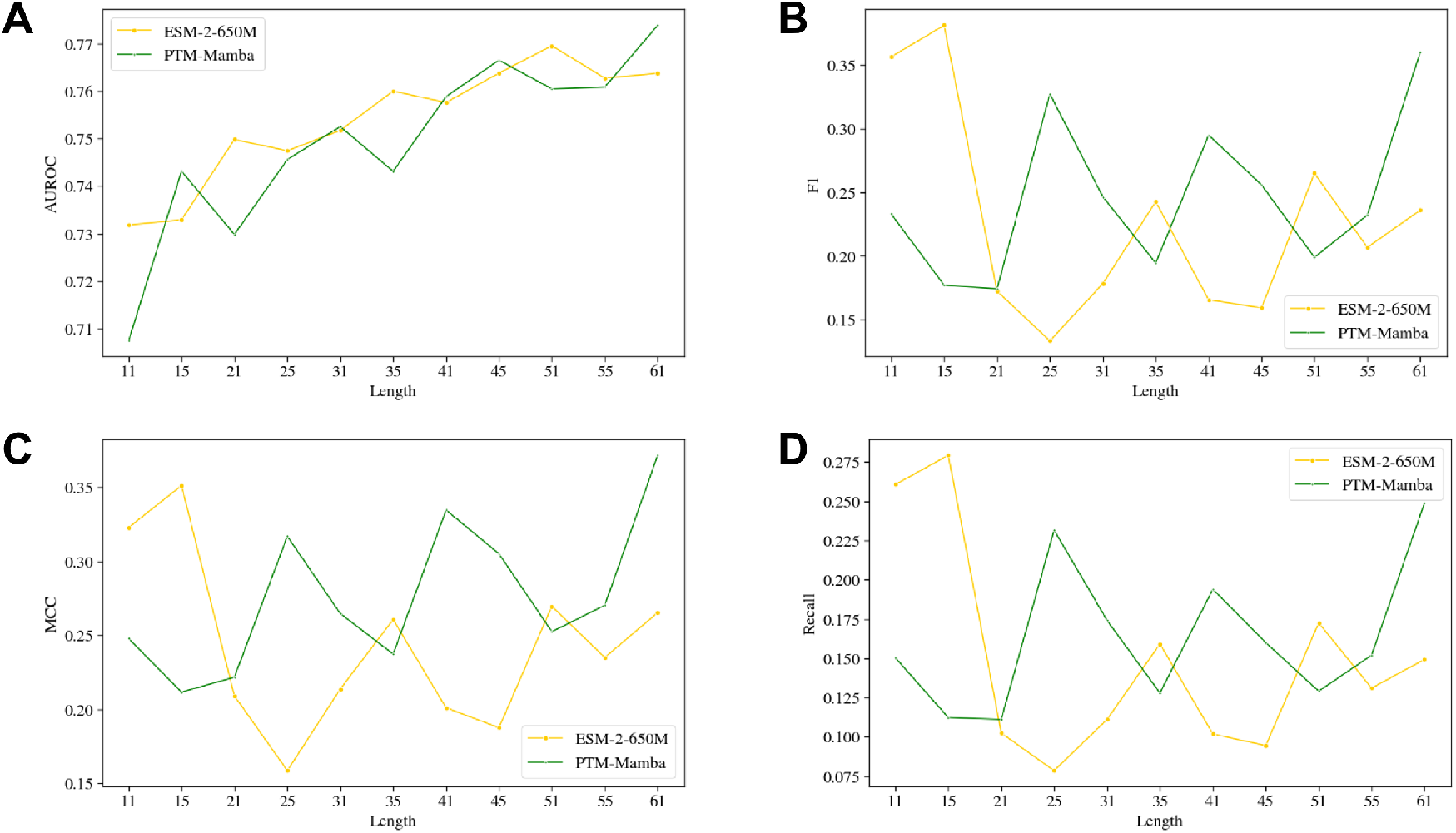
Alternate metrics of non-histone acetylation site prediction. A) AUROC of site prediction for different length input protein sequences. B) F1 score of site prediction for different length input protein sequences. C) MCC metric of site prediction for different length input protein sequences. D) Recall of site prediction for different length input protein sequences.

**Supplementary Figure 4:**
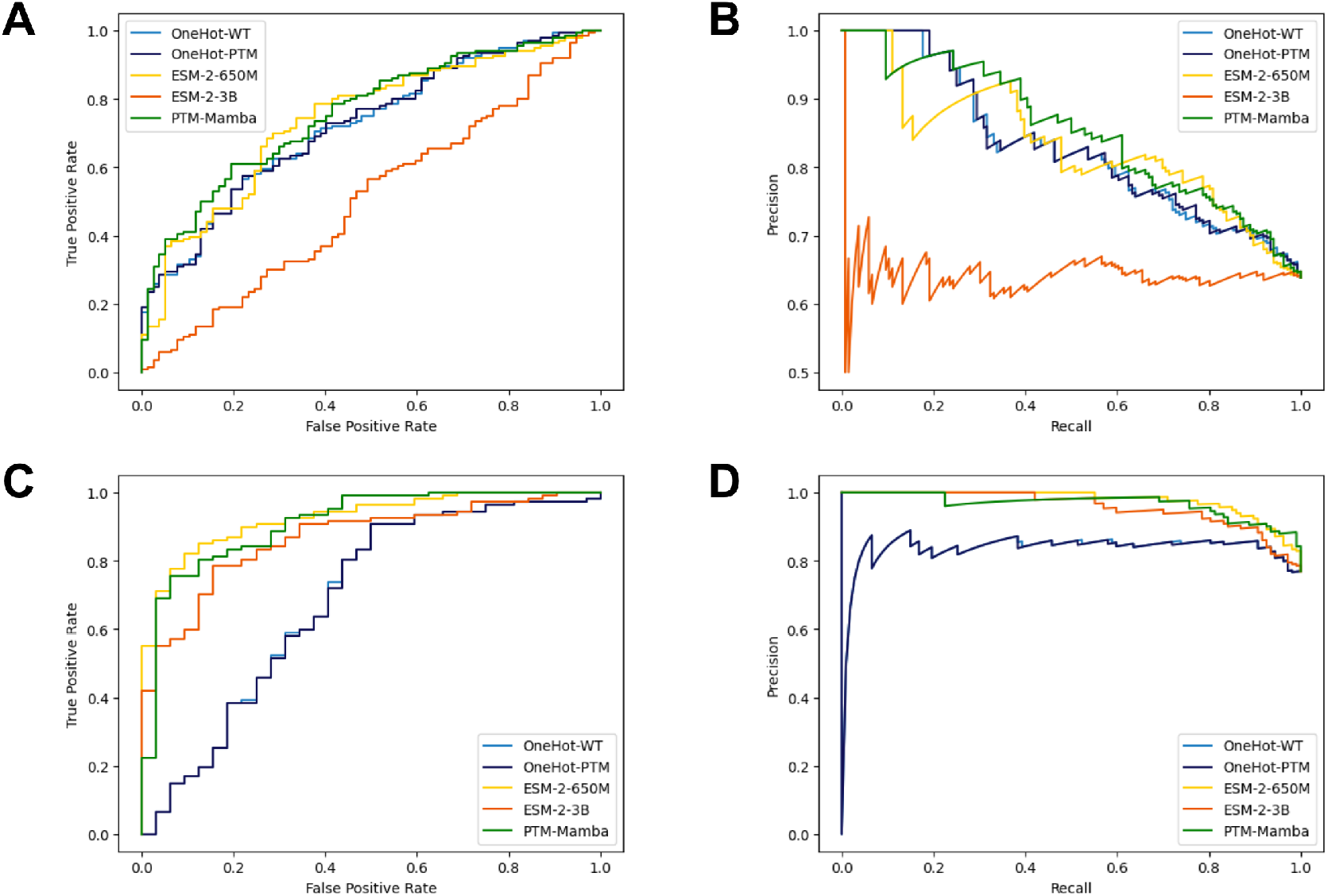
ROC and PRC plots for classification benchmarking. A) Receiver operating characteristic (ROC) for disease association prediction. B) Precision-recall curve (PRC) for disease association prediction. C) ROC for druggability prediction. D) PRC for druggability prediction.

